# Structural and functional characterisation of the interaction between the influenza A virus RNA polymerase and the CTD of host RNA Polymerase II

**DOI:** 10.1101/2023.11.29.569222

**Authors:** Jeremy Keown, Alaa Baazaoui, Marek Šebesta, Richard Štefl, Loic Carrique, Ervin Fodor, Jonathan Grimes

**Affiliations:** Division of Structural Biology, Wellcome Centre for Human Genetics, University of Oxford, Oxford, United Kingdom; Sir William Dunn School of Pathology, University of Oxford, Oxford, United Kingdom; CEITEC–Central European Institute of Technology, Masaryk University, Brno, Czechia; National Centre for Biomolecular Research, Faculty of Science, Masaryk University, Brno, Czechia; Current address: School of Life Sciences, University of Warwick, Coventry, United Kingdom

## Abstract

Influenza A viruses (IAV), causing seasonal epidemics and occasional pandemics, rely on interactions with host proteins for their RNA genome transcription and replication. The viral RNA polymerase utilizes host RNA polymerase II (Pol II) and interacts with the serine 5 phosphorylated (pS5) C-terminal domain (CTD) of Pol II to initiate transcription. Our study, using single-particle electron cryomicroscopy (cryo-EM), reveals the structure of the 1918 pandemic IAV polymerase bound to a synthetic pS5 CTD heptad repeat peptide. The structure shows that the CTD peptide binds at the C-terminal domain of the PA viral polymerase subunit (PA-C) and reveals a previously unobserved position of the 627 domain of the PB2 subunit near the CTD. We identify crucial residues of the CTD peptide mediating interactions with positively charged cavities on PA-C, explaining the preference of the viral polymerase for pS5 CTD. Functional analysis of mutants targeting the CTD-binding site within PA-C reveals reduced transcriptional function with normal replication, while other mutants display defects in both transcription and replication, highlighting the multifunctional role of PA-C in viral RNA synthesis. Our study provides insights into the structural and functional aspects of the influenza virus polymerase-host Pol II interaction and identifies a target for antiviral development.

## INTRODUCTION

Influenza viruses are negative-strand RNA viruses with a genome consisting of eight viral RNA (vRNA) segments organised into viral ribonucleoprotein (vRNP) complexes (1). Each vRNP complex packages one viral genome segment using viral nucleoprotein (NP) and viral RNA-dependent RNA polymerase (RdRp), a heterotrimeric protein, that consists of the polymerase acidic (PA), polymerase basic 1 (PB1) and PB2 subunits (2). During viral infection, the vRNPs are trafficked to the cell nucleus where the viral polymerase performs both viral transcription and replication. Replication, a two-step process, is primer independent and requires de novo initiation by the viral polymerase to synthesise new vRNA, using positive-sense complementary RNA (cRNA) as an intermediate template (2,3). In contrast, viral transcription depends on the interaction of the viral polymerase with the host transcriptional machinery to produce viral positive-sense mRNA. The synthesised viral mRNA contains a 5′ terminal N7- methyl guanosine (m7G) cap and a 3′ polyA tail which makes viral mRNA structurally identical to the host mRNA and allows the virus to hijack the host translation machinery for viral protein synthesis in the cytoplasm (2,4). In a process called ‘cap snatching’, the viral polymerase steals 10-15 nucleotides long capped RNA fragments from nascent capped host RNA, generated by the host RNA polymerase II (Pol II), to prime its own transcription using the PB2 cap-binding and PA endonuclease domains (4,5). Cap snatching takes place during the early stages of Pol II-mediated transcription which enables the viral polymerase to secure access to host RNA caps before being bound by host cap-binding proteins (6). The interaction of the viral polymerase with Pol II has detrimental effects on host transcription and results in inhibition of cellular gene expression (‘host shut-off’) and Pol II degradation at later stages of infection (7–9).

To access caps of nascent host RNA the viral polymerase physically associates with host Pol II. Functional and structural studies confirmed that the viral polymerase selectively recognises the serine 5 phosphorylated (pS5) CTD heptad repeat sequence YSPTpSPS, the signature for initiating Pol II involved in capping (4,10). In structural studies, two distinct binding sites of the CTD to the viral polymerase were observed using polymerase from influenza virus types A, B, and C and synthetic four-repeat pS5 CTD peptides (11–13). Structures of the CTD-bound polymerase of different influenza types revealed shared and distinct binding features between each type, while binding of the Pol II CTD beyond the identified binding sites remains largely unknown. Considering that mammalian Pol II CTD contains 52 heptad repeats of the consensus sequence Y_1_S_2_P_3_T_4_S_5_P_6_S_7_, which far exceeds the length of CTD mimetic peptides, it appears likely that further interactions between the viral RdRP and Pol II CTD take place beyond the identified binding sites (4,10). In addition, it has been reported that the interaction of the viral polymerase with Pol II involves not only the CTD of the large subunit, RPB1, but also other subunits of the 12-subunit holoenzyme such as RPB4 (11,14).

Here, we present the structures of the CTD-bound polymerase of the 1918 pandemic influenza A virus. In these structures, we observe a continuous stretch of the CTD binding site, encompassing three heptad repeats (repeats a-c), on the C-terminal domain of the PA subunit (PA-C). Additionally, we observe a minor population of particles with a unique arrangement of the PB2 C-terminal (PB2-C) region, likely representing a transcriptase intermediate. Based on these structures, we performed mutagenesis of residues in the CTD binding site that are conserved across influenza A virus strains, with the aim of determining their effect on viral polymerase activity using a cell-based vRNP reconstitution assay. Our findings demonstrate that mutagenesis of conserved residues in the CTD binding site results in transcriptional and replicational defects, indicating a potential overlap of binding sites for host factors that facilitate replication and transcription, respectively.

## MATERIALS AND METHODS

### Protein expression and purification

The three polymerase genes from influenza A/Brevig Mission/1/1918 (H1N1) virus were codon optimised for insect cells and synthesised (Synbio Technologies). Genes were then cloned into the Multibac system (15) with protein expression and purification carried out as previously described (16). Cryo-EM studies in this manuscript were carried out using a polymerase that contained a PA D108A endonuclease mutation to reduce RNA degradation. Briefly, Sf9 insect cells were infected with the baculovirus encoding the three viral polymerase genes and harvested 72 hours post infection. One litre of cell pellet was resuspended in 50 mL buffer containing 50 mM HEPES-NaOH, pH 7.5, 500 mM NaCl, 10 % v/v glycerol, 0.05 % w/v n-Octyl beta-D-thioglucopyranoside, 1 mM dithiothreitol which was further supplemented with 1 protease inhibitor cocktail tablet (Roche) and 5 mg RNase A. The cells were lysed with sonication and clarified with centrifugation. The resulting lysate was incubated with IgG Sepharose for three hours before washing the resin with 50 mM HEPES-NaOH, pH 7.5, 500 mM NaCl, 10 % v/v glycerol, 0.05 % w/v n-Octyl beta-D-thioglucopyranoside, 1 mM dithiothreitol. The protein was eluted by the addition of Tobacco Etch Virus (TEV) protease overnight. The eluted protein was concentrated and applied to a Superdex increase S200 10/300 (GE Healthcare) size exclusion column equilibrated with 25 mM HEPES-NaOH, pH 7.5, 500 mM NaCl, 5 % v/v glycerol, 0.5 mM dithiothreitol. Fractions containing pure protein were concentrated and stored at -80 °C.

### Cryo-EM sample preparation

An aliquot of purified polymerase of the A/Brevig Mission/1/1918 (H1N1) influenza virus with a D108A mutation in the PA subunit was defrosted and viral RNA promoters (5′ vRNA 5′- AGUAGAAACAAGGCC-3′, 3′ vRNA 5′-GGCCUGCUUUUGCU*AUU*-3′ with a 3-nucleotide long extension at the 3′ end (italics)) were added to a 1.2 molar excess and incubated on ice for 20 minutes. The sample was then further purified by size exclusion chromatography into a buffer containing 25 mM HEPES-NaOH, pH 7.5, and 500 mM NaCl before being concentrated to 1 mg/ml. A capped RNA primer (5′ m7GpppGAAUGCUAUAAUAGC), with six complementary bases at the 3′ to the 3′ vRNA promoter, was added to this sample to a final concentration of 0.2 mM. Additionally, a pS5 Pol II CTD mimic peptide containing four repeats of the heptapeptide consensus sequence (Y_1_S_2_P_3_T_4_pS_5_P_6_S_7_)_4_, with C-terminal amidation, N-terminal biotinylation and a nine-atom polyethylene glycol spacer between the biotin moiety and the first amino acid (PeptideSynthetics, Peptide Protein Research Ltd) (Supplementary Table 1), was added to a final concentration of 0.2 mM. Immediately prior to grid preparation the sample with diluted 1:3 with a buffer containing HEPES-NaOH, pH 7.5, 37.5 mM NaSCN and 0.0075 % (v/v) Tween20. A volume of 3.5 μL of the sample was applied to a freshly glow-discharged Quantifoil Holey Carbon R2/1, 200 mesh copper grid which was blotted for 3.5 seconds and plunge frozen in liquid ethane. All grids were prepared using a Vitrobot mark VI (FEI) at 100 % humidity and 20 °C.

### Cryo-EM image collection and processing

Data were collected on a 300 kV Titan Krios with a K2 Summit camera (Gatan) and a GIF Quantum energy filter at the Oxford Particle Imaging Centre. Data were collected using SerialEM (17). Figures were prepared using ChimeraX (18). Data processing is graphically summarised in Supplementary Figure 1 and data collection and processing details are presented in Supplementary Table 2. cryoSPARC V4-4.2 (19,20) was to perform patch motion correction, patch CTF estimation, and picking of initial particles using the blob picker. Micrographs with poor pre-processing statistics were manually removed. After 2D classification and generation of an initial high resolution consensus refinement these particles were used to train a topaz model (21) and repick particles in the dataset. Particles were combined and duplicates removed.

After performing rounds of 3D classification on the CTD peptide dataset, a consensus refinement involving 303,608 particles resulted in a map with a resolution of 3.22 Å. A model was built into this reconstruction using the 7NHX as a starting model. The model was manually built in COOT before refinement in real space using PHENIX (22). Further 3D classification yielded a reconstruction which contained extra density corresponding to PB2-C. To aid interpretation of the density, the map was modified using deepEMhancer (23). In this modified map, the previously mentioned model, encompassing the PA, PB1, and N-terminal region of PB2 (PB2-N) subunits, was positioned. Subsequently, the PB2-C domains (extracted from PDB 7NHX) were placed into the corresponding density. Due to the low resolution of the density in this region, the domains were manually connected in COOT before restrained rigid-body real-space and ADP refinement.

### Cells and plasmids

Human embryonic kidney (HEK) 293T cells (293T) were grown in Dulbecco’s modified Eagle’s medium (DMEM, Gibco) supplemented with 10 % fetal bovine serum (FBS, Gibco) and cultured at 37 °C with 5 % CO_2_. The pCAGGS-PA-1918, pCAGGS-PB1-1918, pCAGGS- PB2-1918, pCAGGS-NP-1918 plasmids expressing the vRNP components of influenza A/Brevig Mission/1/1918 virus, have been described previously (24). The pPOLI-NA-RT plasmid expressing a segment 6 vRNA template under the control of a human RNA polymerase I (Pol I) has also been described (25). Mutations in pCAGGS-PA-1918 were introduced using site-directed mutagenesis. All constructs were confirmed by Sanger sequencing.

### vRNP reconstitution assay, RNA isolation and analysis

To compare the activity of mutant viral polymerases in cell culture, HEK 293T cells were grown to 80 % confluency in six-well dishes and transfected with 0.5 μg of each of wild-type (WT) or mutant pCAGGS-PA-1918, pCAGGS-PB1-1918, pCAGGS-PB2-1918, pCAGGS-NP-1918, and pPOLI-NA-RT plasmids. For the negative control, pCAGGS-PA-1918 was replaced with an empty vector. Transfections were performed using Lipofectamine 2000 (Invitrogen) according to the manufacturer’s instructions. Total RNA from transfected cells was extracted using TRI reagent (Sigma) according to the manufacturer’s instructions 24 h post transfection. Levels of mRNA, cRNA, and vRNA were analysed by primer extension assay as previously described (26). In brief, extracted RNA was reverse transcribed by SuperScript III reverse transcriptase (Invitrogen) and ^32^P-labelled NA-specific primers along with a primer targeting endogenous 5S rRNA as an internal control. Primer extension products were separated by 6 % denaturing PAGE with 7 M urea in TBE buffer and bands were detected by phosphorimaging on an FLA-5000 scanner (Fuji). The cDNA was analysed using ImageJ (Fiji) and Prism 8 (GraphPad).

### Immunoblotting

Immunoblotting was performed to determine expression levels of WT and mutant PA in HEK293T cells. PA, NP (transfection control) and GAPDH (loading control) were probed overnight at 4 °C with rabbit anti-PA (6) (1:500), rabbit anti-NP (1:5000) (Genetex), and rabbit anti-GAPDH (1:1000) (Cell Signalling Technology). Goat anti-rabbit antibody conjugated to horseradish peroxidase (HRP) was used as secondary antibody (1:10,000) (Genetex). Bands were detected using Amersham ECL Western blotting detection reagents (GE Healthcare).

### Design and synthesis of Pol II CTD mimic peptides

Peptides were chemically synthesized by solid-phase peptide synthesis (Peptide Protein Research Ltd.). The designed peptides contain four repeats of the heptapeptide consensus sequence of the Pol II CTD (YSPTSPS) with modifications to mimic different phosphorylation states of serine residues at position 2, position 5, and position 7 of the CTD. Full amino acid sequences are detailed in (Supplementary Table 1). All peptides were synthesized with C-terminal amidation, N-terminal biotinylation, and included a nine-atom polyethylene glycol spacer between the biotin moiety and the first amino acid. Peptides were purified by high-performance liquid chromatography to at least 90% purity and peptide quality was confirmed by mass spectrometry.

### Binding assay of influenza virus polymerase to Pol II CTD

The binding of purified viral polymerase to synthetic four-heptad repeat Pol II CTD mimic peptides (Supplementary Table 1) was performed as previously described (27,28). In brief, biotinylated Pol II CTD mimic peptide (20 µg) were immobilised on 10 µl streptavidin agarose resin (Thermo Scientific) for 2 hours at 4 °C in binding buffer (10 mM HEPES [PAA catalogue no. S11-001], 150 mM NaCl, 0.1 % (v/v) Igepal, 1× Halt protease inhibitor cocktail [Pierce], 1% (w/v) bovine serum albumin (BSA)), followed by washing with wash buffer (10 mM HEPES [PAA catalogue no. S11-001], 150 mM NaCl, 0.1 % (v/v) Igepal, 1 mM PMSF). After washing of the beads with wash buffer, 4 µg of 1918 influenza virus polymerase in binding buffer were incubated with the peptide-bound beads for 2 hours at 4 °C. Beads were washed and boiled in SDS-PAGE sample buffer at 95 °C for 5 minutes, followed by SDS-PAGE analysis and silver staining to visualise protein bands according to the manufacturer’s instructions (Invitrogen).

## RESULTS

### Effect of alternative phosphorylation patterns of the Pol II CTD on binding the influenza A virus polymerase

The Pol II CTD exhibits remarkable functional flexibility because many of the amino acids in the heptapeptide can undergo posttranslational modifications and various combinations of these covalent marks are selectively recognized by different protein partners (29–31). Of the amino acid residues composing the consensus heptad, serines at positions 2, 5 and 7 (abbreviated as S2, S5 and S7, respectively) as well as the tyrosine at position 1 (Y1) and the threonine at position (T4), can be phosphorylated, with S2 and S5 phosphorylations being the most prominent posttranslational modifications. The influenza virus RNA polymerase has been shown to have a strong preference for the pS5 CTD over pS2 and unphosphorylated CTD. However, combinations of these phosphorylation patterns and the effect of pS7 have not yet been investigated.

To further characterise the binding preference of the influenza virus polymerase to Pol II CTD, we used biotinylated Pol II CTD mimic peptides containing four repeats (designated repeats a-d) of the conserved heptapeptide repeat (Y_1_S_2_P_3_T_4_pS_5_P_6_S_7_) with a combination of phosphorylation patterns (Figure 1, Supplementary Table 1). We found that the polymerase of the 1918 pandemic influenza virus bound to the pS5 peptide and we also detected low but above background levels of binding to pS7. Interestingly, none of the other peptides were bound by the viral polymerase above background as determined by polymerase binding to a peptide with a scrambled sequence. Overall, this experiment shows that the influenza virus polymerase has a striking preference for pS5 peptide compared to pS2 and pS7 and additional phosphorylations in the pS5 peptide interfere with binding. This binding preference clearly links the influenza virus polymerase to the initiating form of Pol II engaged in capping its nascent RNA (32).

**Figure 1.**
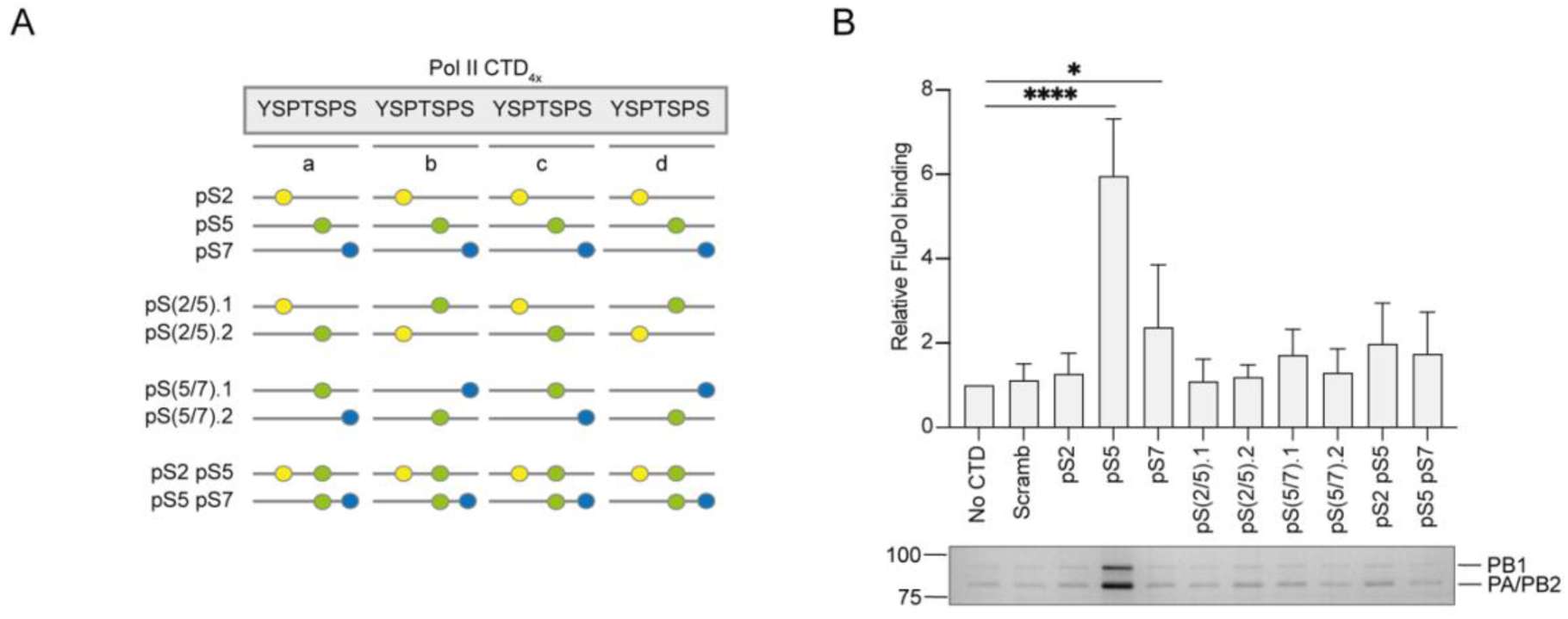
Effect of serine phosphorylations of Pol II CTD on binding the 1918 pandemic H1N1 influenza A virus polymerase. (**A**) Schematic of peptides used in the binding assay with the four heptad repeats (designated repeats a, b, c, and d) depicted as grey lines. Serine phosphorylations (pS) are shown as coloured circles (pS2 (yellow), pS5 (green), pS7 (blue)). (**B**) Binding of the influenza virus polymerase to serine phosphorylated Pol II CTD mimic peptides. Top panel, quantification of influenza virus polymerase (FluPol) binding from n = 3 independent binding assays. Data are mean ± s.e.m. Ordinary one-way ANOVA was used to compare the relative polymerase binding in the presence and absence of CTD. *, p<0.05; ****, p<0.0001. Bottom panel, a representative silver-stained gel.

### Structure of the 1918 pandemic influenza A virus polymerase bound to Pol II CTD mimic peptide

To characterise the binding of the CTD of host Pol II to the 1918 pandemic influenza A virus polymerase, we used recombinant influenza virus polymerase expressed in insect cells and a four-repeat synthetic pS5 Pol II CTD mimic peptide. Cryo-EM analysis of the complex generated a high-quality map to a global resolution of 3.22 Å (Figure 2A) comprising the polymerase core (PA-C, PB1, PB2-N domains), the PA-C endonuclease domain, 15 bases of the 5′ vRNA promoter, eight bases from the 5′ end of the 3′ vRNA promoter, and continuous density for 25 amino acids of the CTD peptide (Figure 2B). Though capped RNA was added to the complex this was not observed either in the RdRp active site or the PB2 cap-binding domain.

**Figure 2.**
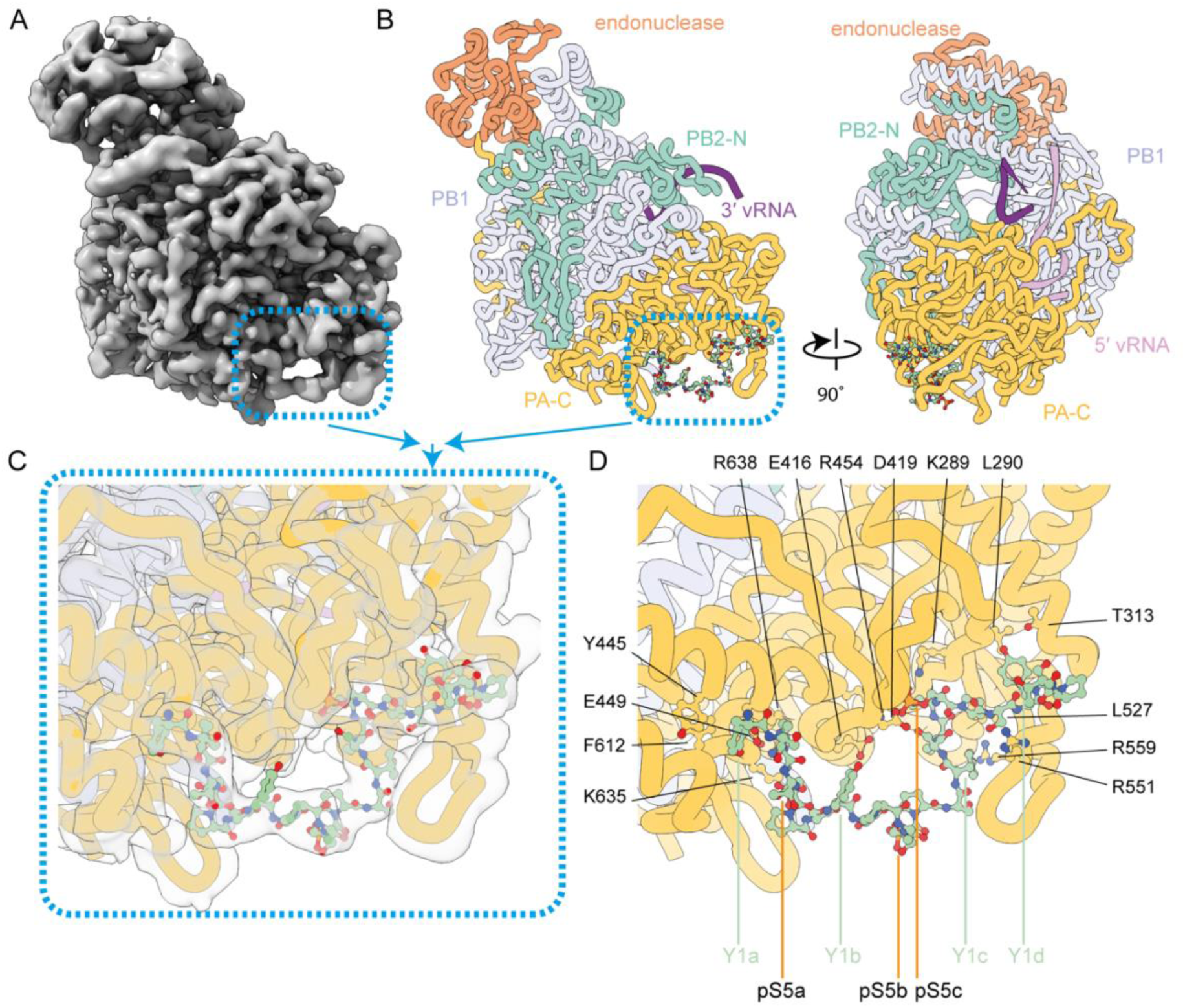
Structure of the 1918 pandemic H1N1 influenza A virus polymerase bound to a pS5 Pol II CTD mimic peptide. (**A-B**) deepEMhancer modified Cryo-EM map (A) and model in two orientations showing the position of the CTD peptide bound at PA-C domain (B). (**C-D**) Close-up views detailing the interaction of the CTD peptide with the PA-C domain showing the fit of the model (C) and amino acids involved in mediating the interactions (D).

The PB2-C domains were not observed in the consensus refinement nor were the 10 bases from the 3′ end of the promoter vRNA. The global polymerase conformation is that of a transcriptase as characterised by the position of the endonuclease domain packing against the PB1 C-terminal helices (33–35). This endonuclease conformation is distinct from the arrangement of the endonuclease domain in any of the previously observed structures (36). The CTD occupies an elongated binding site along the PA-C domain, where we observe 25 of the 28 residues from the peptide (Figure 2C). The phosphorylated peptide buries a total surface area of 1160 Å^2^ with the interaction mediated predominantly by the first (designated repeat a), third (designated repeat c), and the N-terminal half of the fourth heptad repeat (designated repeat d). The phosphorylated serine residues pS5a and pS5c of the peptide are highly ordered and are coordinated into binding sites formed by K635/R638 and K289/R454, respectively. E416 and E457 aid in the correct positioning of R454 to coordinate pS5c. Other amino acid residues of the polymerase interact with unmodified residues from the CTD peptide. Specifically, residues Y445, E449, and F612 interact with Y1a and D419 may contact Y1b. At the C-terminal end of the resolved part of the peptide L290 and Y313 contact Y1d, L527 may interact with P6c, and R551/R559 contact the peptide as it leaves the surface of the PA-C over the top of the PA 550- loop. The phosphorylated pS5b was more poorly ordered and does not appear to contact the polymerase core (Figure 2D). All residues interacting with the peptide are from the PA subunit.

### Arrangement of PB2-C in the Pol II CTD-bound influenza virus polymerase

While most particles in our cryo-EM dataset lacked density for the flexible PB2-C domains, including the cap-binding domain, mid-link, 627 and nuclear localisation signal (NLS) domains, approximately 5.3 % of the particles from the consensus refinement contained additional electron density, suggesting that in these particles the PB2-C domains had become ordered (Figure 3A, Supplementary Figure 1). By aligning the core of previously determined polymerase structures we were able to accurately position the cap-binding domain and mid-link domain in the density (Figure 3B). The position of these two domains was essentially identical to that of the recently determined transcription intermediate structure (PDB ID 7NHX) (27). In this model the mid-link domain has remained in close contact with the core of the polymerase (in a position common to transcriptase conformations) while the cap-binding domain has retracted away from the core and primer entry channel. A putative role of this conformation was recently proposed, as an intermediate state between transcription and replication (37). To aid our attempts to unambiguously orient the 627 domain and/or the NLS we utilised deepEMhancer to modify the map. From this modified map we could correctly position the 627 domain such that the connectivity to the mid-link was maintained and with a good fit to the modified density. No alternate orientation of the domain was possible.

**Figure 3.**
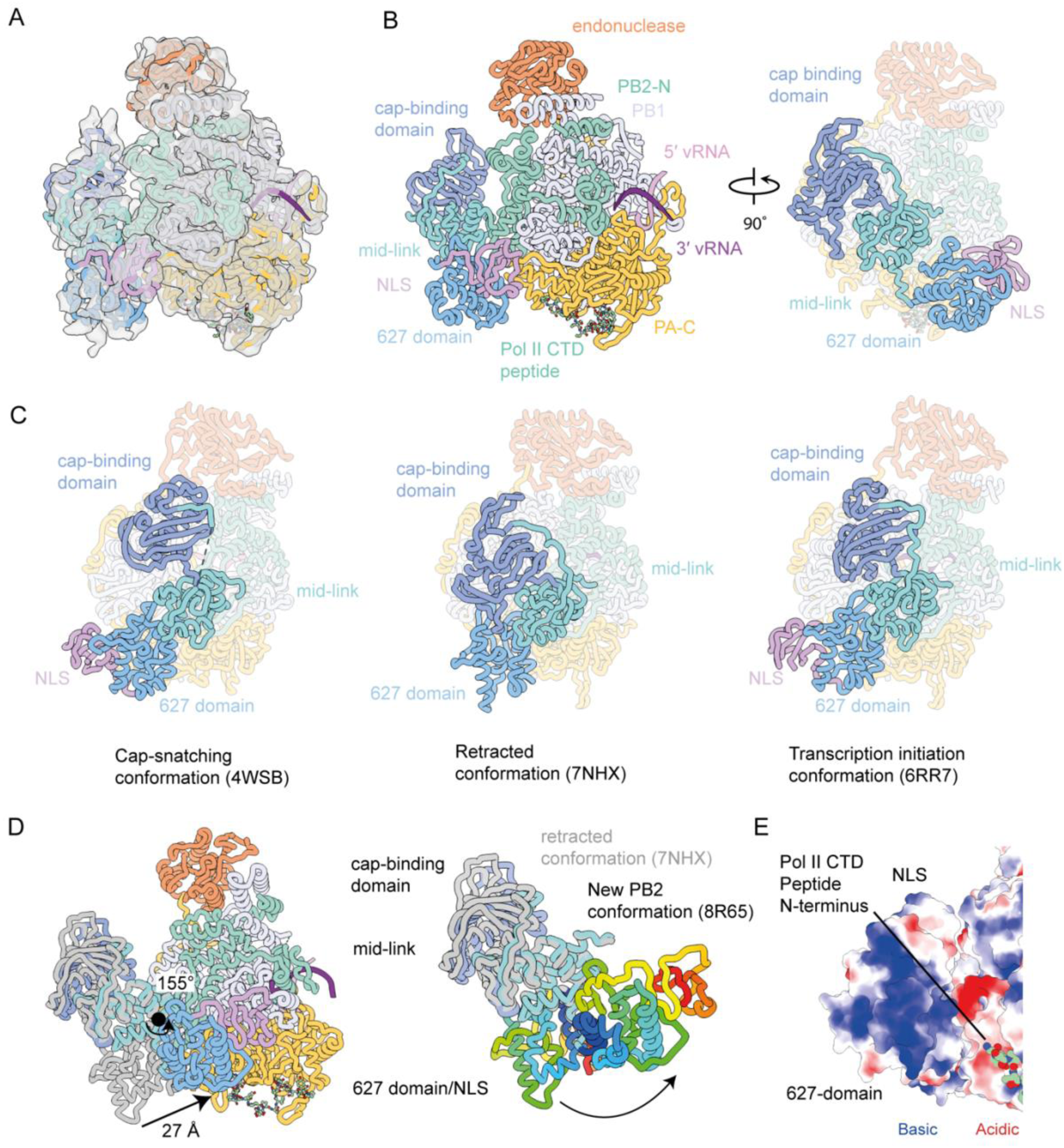
Structure of the 1918 pandemic H1N1 influenza A virus polymerase bound to a pS5 Pol II CTD mimic peptide with resolved PB2-C domains. (**A-B**) deepEMhancer modified Cryo-EM map (A) and model in two orientations showing the domain structure with the arrangement of the PB2-C domains highlighted on the right (B). (**C**) Previously determined structures of the influenza virus polymerase with different arrangements of the PB2-C domains highlighted. (**D**) Model of the CTD-bound polymerase showing the movement of the rainbow coloured PB2 627 domain while transitioning from the retracted conformation to that observed in the CTD-bound structure. (**E**) Electrostatic analysis of the PB2 627 domain that is oriented towards the N-terminus of the CTD peptide shows a highly basic surface.

The 627 domain and NLS in this model adopts a position that is distinct from its location in either a replicating polymerase, encapsidating polymerase, or previously observed transcription initiation or cap-snatching structures (Figure 3C) (36). In this model the 627 domain and NLS has rotated 155° and moved 27 Å from the transcription initiation position (Figure 3D). The movement changes the interaction with the polymerase from contacting the PA and PB1 subunits through a three stranded beta-sheet formed by residues 640-676 such that it now contacts the PA-C and PB2-N2 domains through residues 643-654. This interface is smaller than observed in the transcription initiation conformation resulting in this domain being poorly resolved. A small interface was formed between the NLS and residues from the PB1 (583–587), PB2 (125–127), and PA (430–432) subunits. Due to the low resolution, we were unable to accurately describe residue contacts in these regions and will instead limit our analysis to the region of the domain. The relocation of the 627 domain and NLS brings it proximal to the N-terminus of the Pol II CTD peptide.

We do not observe any additional peptide density in this reconstruction that would suggest a direct contact between the CTD and the PB2-C. However, given the low-resolution maps, we cannot exclude the possibility of a direct contact. In fact, electrostatic analysis of the surface of the 627 domain that is oriented towards the N-terminus of the CTD peptide shows a highly basic surface (Figure 3E). This surface would likely be receptive to binding of the negatively charged phosphorylated CTD peptide. Recent structural evidence of Pol II CTD binding to the influenza B virus polymerase demonstrates a large additional CTD binding site across the surface of the 627 domain (11,13). However, corresponding residues from the influenza B polymerase are not conserved in influenza A strains. Deletion of the 627 domain from the influenza B virus polymerase dramatically reduced binding to Pol II CTD while a similar deletion did not affect the ability of the influenza A virus polymerase to bind CTD (11). It could be that the loss of binding from the deletion of the 627 domain is compensated by the much larger binding surface of the CTD peptide on the influenza A virus PA-C domain.

### Effect of mutations in the PA-C Pol II CTD binding site on polymerase function

To address the role of the identified Pol II CTD binding site in PA-C in polymerase function, we used a vRNP reconstitution assay followed by the analysis of RNA by primer extension. We co-expressed the three polymerase subunits, nucleoprotein, and segment 6 encoding the neuraminidase vRNA in HEK 293T cells to assemble vRNPs, the minimal viral complex required for transcription (mRNA synthesis) and genome replication (cRNA and vRNA synthesis) (Supplementary Figure 2). Based on our structural model and previously determined RdRP structures from IAV strains, we mutated 14 conserved residues in the PA-C domain that we hypothesised to influence CTD binding (Supplementary Figure 3) (11,13,27,35). Only low levels of vRNA, expressed from the transfected plasmid, were observed if the PA subunit was omitted from the transfection (Figure 4). However, in the presence of a complete wild-type polymerase, we observed mRNA and cRNA, as well as a significant increase in vRNA levels, indicating that the viral polymerase transcribes and replicates the input vRNA. Comparing the RNA levels produced by the wild-type and mutant polymerases, we found that mutations K289A, Y445A, E449A, R454A and K635A affected transcription with relatively little if any effect on replication. The phenotype of these mutants is fully consistent with CTD binding being required for viral transcription, as this interaction facilitates access of the viral polymerase to capped RNA fragments generated from nascent Pol II transcripts. Other mutations such as L290A, T313A, D419A, L527A, R559A, F612A and T639A had no or only a small effect on transcription indicating that these amino acid residues do not contribute critical interactions to CTD binding. Surprisingly, two mutations, E416A and R551A specifically reduced cRNA synthesis, suggesting that amino acid residues around the Pol II CTD binding site also participate in the replication of the viral RNA genome.

**Figure 4.**
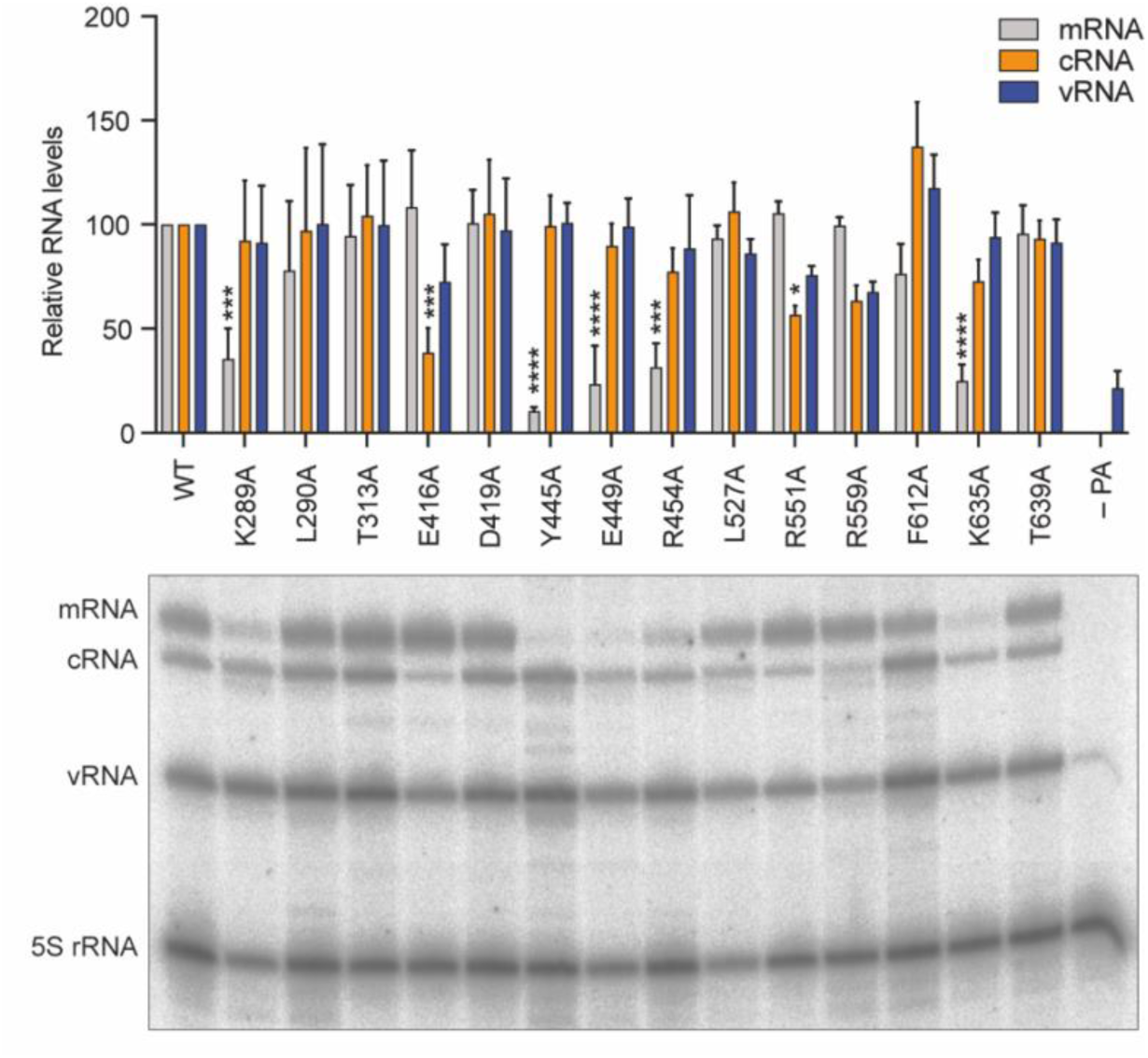
The effect of mutations in the CTD binding site of the 1918 pandemic H1N1 influenza A virus polymerase on viral RNA synthesis. HEK 293T cells were transfected to express the 1918 influenza virus polymerase subunits PA (WT and mutant), PB1, and PB2, as well as segment 6 vRNA. For the negative control, the plasmid expressing PA was replaced with an empty vector. Total RNA was isolated 24 hours post-transfection and was analysed by primer extension assay. Top panel, quantification of mRNA, cRNA, and vRNA levels from n=3 independent vRNP reconstitution assays. Bottom panel, a representative primer extension analysis. 5S rRNA was used as a loading control. The mean signal intensity is shown relative to the signal intensity from WT 1918 viral polymerase. Data are mean ± s.e.m. Ordinary two- way ANOVA was employed to determine significant differences with *p < 0.05; ***p < 0.001; and ****p < 0.0001.

## DISCUSSION

In this study we have solved the solution state structure of the 1918 pandemic H1N1 influenza virus polymerase bound to pS5 Pol II CTD using cryo-EM. The structure revealed a continuous density for the CTD peptide with 25 out of the 28 amino acid residues fully resolved. The CTD peptide was found to bind exclusively at the PA-C with two of the phosphorylated serine residues playing prominent roles in guiding the interaction by being accommodated in highly basic grooves. The interaction interface overlaps with that previously observed in a study using polymerase of a bat H17N10 influenza A virus (13). However, in contrast to this study that observed two separate binding sites for the CTD here we identify a continuous density with the CTD linking the two binding sites. Our structure is the first IAV polymerase structure determined by cryo-EM, revealing that binding of CTD in solution does not stabilise the cap-snatching or transcription initiation complexes and suggesting that a longer CTD peptide and/or other interacting partners are required. While in most particles the PB2-C domains remained unstructured, we observed in a small population the PB2 cap-binding, mid-link, 627, and NLS domains in an arrangement not previously observed. Interestingly, in this novel arrangement the 627 domain is positioned close to the N-terminus of the CTD peptide providing a potential basic surface for additional interactions with the acidic CTD. Binding of the CTD across PA- C and the adjacent 627 domain could stabilise the 627 domain and, consequently, enable the PB2 cap-binding and PA endonuclease domains to assume a cap-snatching competent conformation. Using Pol II CTD peptides composed of more than four heptad repeats could reveal further insights into the interaction of the influenza virus polymerase with Pol II CTD.

The Pol II CTD can be phosphorylated at S2, S5, and S7 and on T4 of the heptad repeat (30). Binding assays of the influenza A virus polymerase to Pol II CTD peptides containing different phosphorylation patterns revealed a striking preference for pS5 CTD peptides over pS2 or pS7 peptides. Interestingly, combining pS5 with pS2 or pS7 in various arrangements in the same peptide lead to a dramatic reduction of polymerase binding suggesting that the viral polymerase engages with Pol II at a very precise point in the transcription cycle when the capping enzyme is recruited to Pol II (38,39). Analysis of our structure explains how additional phosphorylation at S2 and S7 would disrupt CTD binding. Phosphorylation at S2a and S2b would likely be well accommodated as these residues are exposed towards the solvent. On the other hand, phosphorylation at S7a, S7b, S7c, and S2d could be accommodated but would likely require some rearrangement of the peptide. Phosphorylation of S2c would be unlikely to be accommodated given the tight contact with the polymerase and the nearby highly negative charge on S5c.

We observed that mutations of PA-C at the CTD binding site cause defects in both transcription and replication. Transcription was affected by mutations in K289A, Y445A, E449A, R454A and K635A of the PA subunit with little to no effects on genome replication. These amino acid residues, along with the previously extensively studied R638 (13,40), are involved in direct interactions with pS5 of the first and third heptad repeat of the Pol II CTD (pS5a and pS5c) as well as Y1 of the first repeat (Y1a), highlighting that interactions with these three amino acids are the most critical for CTD binding. These results are consistent with CTD binding being important for viral mRNA synthesis by enabling cap-snatching. Interestingly, we observed reduced replication of the PA E416A and R551A mutants with little effect on transcription. Recent studies of the influenza C virus polymerase in complex with human and chicken ANP32A revealed that the ANP32A binding site overlaps with part of the Pol II CTD binding site (41) (Supplementary Figure 4). ANP32 proteins have been shown to be essential factors for the genome replication of all three types of influenza viruses (41–43) and influenza A virus likely also interacts with ANP32A through its PA-C domain. We speculate that PA mutations E416A and R551A might inhibit genome replication through interfering with ANP32A binding. Further structural studies using influenza A virus polymerase in complex with ANP32A could provide further insights into how the alternate binding of Pol II CTD and ANP32A at the same site in PA-C regulate transcription and replication by the viral polymerase. The importance of this site in the viral replication cycle has recently been highlighted through the use of nanobodies (27). Specifically, two nanobodies which bind PA- C at sites adjacent to the CTD peptide binding site and have been shown to block CTD binding, inhibited both mRNA and cRNA synthesis and, consequently, viral growth.

In summary, our work has uncovered the structural basis of the binding of the CTD of Pol II to the polymerase of the 1918 pandemic influenza A virus. Future research aiming to understand cap-snatching in molecular detail within the context of mammalian Pol II will require the inclusion of additional accessory proteins suspected to contribute to the complex formation between Pol II and the viral polymerase. The highly repetitive and intrinsically disordered nature of the CTD drives phase separation and leads to the recruitment of Pol II and transcription factors that together form condensates (44–48). In the future, CTD-driven phase separation should be investigated within the context of viral infection to facilitate a spatiotemporal understanding of the links between the host transcriptional apparatus and viral transcription and replication beyond the PA-C.

## DATA AVAILABILITY

Map and models have been deposited in the Electron Microscopy and Protein Data Bank, respectively. The structure of the 1918 pandemic IAV polymerase bound to vRNA and Pol II CTD peptide is deposited under EMD-19845 and PDB-8R60, the structure of the 1918 pandemic IAV polymerase bound to vRNA and Pol II CTD peptide with ordered PB2 is deposited under EMD-18947 and PDB-8R65.

Source data as well as plasmids are available upon request.

## SUPPLEMENTARY DATA

Supplementary Data are available at NAR Online.

## ACKNOWLEDGEMENTS

We thank Haitian Fan and Jane Sharps for their assistance with protein expression and advice on the functional assays. We also thank members of the Grimes and Fodor laboratories for helpful comments and discussions.

## Author contributions

All authors have contributed to the design of the experiments. J.K. performed the structural analysis and data processing. A.B. performed all biochemical experiments. All authors discussed the results and commented on the manuscript with the initial draft prepared by A.B. and J.K.

## REFERENCES

1. Eisfeld, A.J., Neumann, G. and Kawaoka, Y. (2015) At the centre: influenza A virus ribonucleoproteins. Nat Rev Microbiol, 13, 28–41.

2. Te Velthuis, A.J. and Fodor, E. (2016) Influenza virus RNA polymerase: insights into the mechanisms of viral RNA synthesis. Nat Rev Microbiol, 14, 479–493.

3. Zhu, Z., Fodor, E. and Keown, J.R. (2023) A structural understanding of influenza virus genome replication. Trends Microbiol, 31, 308–319.

4. Walker, A.P. and Fodor, E. (2019) Interplay between Influenza Virus and the Host RNA Polymerase II Transcriptional Machinery. Trends Microbiol, 27, 398–407.

5. Wandzik, J.M., Kouba, T. and Cusack, S. (2021) Structure and Function of Influenza Polymerase. Csh Perspect Med, 11.

6. Engelhardt, O.G., Smith, M. and Fodor, E. (2005) Association of the influenza A virus RNA-dependent RNA polymerase with cellular RNA polymerase II. Journal of virology, 79, 5812–5818.

7. Vreede, F.T., Chan, A.Y., Sharps, J. and Fodor, E. (2010) Mechanisms and functional implications of the degradation of host RNA polymerase II in influenza virus infected cells. Virology, 396, 125–134.

8. Bauer, D.L.V., Tellier, M., Martinez-Alonso, M., Nojima, T., Proudfoot, N.J., Murphy, S. and Fodor, E. (2018) Influenza Virus Mounts a Two-Pronged Attack on Host RNA Polymerase II Transcription. Cell Rep, 23, 2119-+.

9. Rodriguez, A., Perez-Gonzalez, A. and Nieto, A. (2007) Influenza virus infection causes specific degradation of the largest subunit of cellular RNA polymerase II. Journal of virology, 81, 5315–5324.

10. Krischuns, T., Lukarska, M., Naffakh, N. and Cusack, S. (2021) Influenza Virus RNA-Dependent RNA Polymerase and the Host Transcriptional Apparatus. Annu Rev Biochem, 90, 321–348.

11. Krischuns, T., Isel, C., Drncova, P., Lukarska, M., Pflug, A., Paisant, S., Navratil, V., Cusack, S. and Naffakh, N. (2022) Type B and type A influenza polymerases have evolved distinct binding interfaces to recruit the RNA polymerase II CTD. PLoS pathogens, 18, e1010328.

12. Serna Martin, I., Hengrung, N., Renner, M., Sharps, J., Martínez-Alonso, M., Masiulis, S., Grimes, J.M. and Fodor, E. (2018) A Mechanism for the Activation of the Influenza Virus Transcriptase. Mol Cell, 70, 1101–1110.e1104.

13. Lukarska, M., Fournier, G., Pflug, A., Resa-Infante, P., Reich, S., Naffakh, N. and Cusack, S. (2017) Structural basis of an essential interaction between influenza polymerase and Pol II CTD. Nature, 541, 117–121.

14. Morel, J., Sedano, L., Lejal, N., Da Costa, B., Batsche, E., Muchardt, C. and Delmas, B. (2022) The Influenza Virus RNA-Polymerase and the Host RNA-Polymerase II: RPB4 Is Targeted by a PB2 Domain That Is Involved in Viral Transcription. Viruses, 14.

15. Bieniossek, C., Imasaki, T., Takagi, Y. and Berger, I. (2012) MultiBac: expanding the research toolbox for multiprotein complexes. Trends Biochem Sci, 37, 49–57.

16. York, A., Hengrung, N., Vreede, F.T., Huiskonen, J.T. and Fodor, E. (2013) Isolation and characterization of the positive-sense replicative intermediate of a negative-strand RNA virus. Proceedings of the National Academy of Sciences of the United States of America, 110, E4238–E4245.

17. Mastronarde, D.N. (2003) SerialEM: A Program for Automated Tilt Series Acquisition on Tecnai Microscopes Using Prediction of Specimen Position. Microscopy and Microanalysis, 9, 1182–1183.

18. Pettersen, E.F., Goddard, T.D., Huang, C.C., Meng, E.C., Couch, G.S., Croll, T.I., Morris, J.H. and Ferrin, T.E. (2021) UCSF ChimeraX: Structure visualization for researchers, educators, and developers. Protein Sci, 30, 70–82.

19. Punjani, A., Rubinstein, J.L., Fleet, D.J. and Brubaker, M.A. (2017) cryoSPARC: algorithms for rapid unsupervised cryo-EM structure determination. Nat Methods, 14, 290–296.

20. Punjani, A., Zhang, H. and Fleet, D.J. (2020) Non-uniform refinement: adaptive regularization improves single-particle cryo-EM reconstruction. Nature Methods, 17, 1214–1221.

21. Bepler, T., Morin, A., Rapp, M., Brasch, J., Shapiro, L., Noble, A.J. and Berger, B. (2019) Positive-unlabeled convolutional neural networks for particle picking in cryo-electron micrographs. Nature Methods, 16, 1153–1160.

22. Afonine, P.V., Poon, B.K., Read, R.J., Sobolev, O.V., Terwilliger, T.C., Urzhumtsev, A. and Adams, P.D. (2018) Real-space refinement in PHENIX for cryo-EM and crystallography. Acta Crystallogr D Struct Biol, 74, 531–544.

23. Sanchez-Garcia, R., Gomez-Blanco, J., Cuervo, A., Carazo, J.M., Sorzano, C.O.S. and Vargas, J. (2021) DeepEMhancer: a deep learning solution for cryo-EM volume post-processing. Communications Biology, 4, 874.

24. Tumpey, T.M., Basler, C.F., Aguilar, P.V., Zeng, H., Solórzano, A., Swayne, D.E., Cox, N.J., Katz, J.M., Taubenberger, J.K., Palese, P. et al. (2005) Characterization of the reconstructed 1918 Spanish influenza pandemic virus. Science, 310, 77–80.

25. Fodor, E., Devenish, L., Engelhardt, O.G., Palese, P., Brownlee, G.G. and Garcia-Sastre, A. (1999) Rescue of influenza A virus from recombinant DNA. Journal of virology, 73, 9679–9682.

26. te Velthuis, A.J.W., Robb, N.C., Kapanidis, A.N. and Fodor, E. (2016) The role of the priming loop in influenza A virus RNA synthesis. Nature Microbiology, 1, 16029.

27. Keown, J.R., Zhu, Z., Carrique, L., Fan, H., Walker, A.P., Serna Martin, I., Pardon, E., Steyaert, J., Fodor, E. and Grimes, J.M. (2022) Mapping inhibitory sites on the RNA polymerase of the 1918 pandemic influenza virus using nanobodies. Nature communications, 13, 251.

28. Martínez-Alonso, M., Hengrung, N. and Fodor, E. (2016) RNA-Free and Ribonucleoprotein-Associated Influenza Virus Polymerases Directly Bind the Serine- 5-Phosphorylated Carboxyl-Terminal Domain of Host RNA Polymerase II. J Virol, 90, 6014–6021.

29. Dieci, G. (2021) Removing quote marks from the RNA polymerase II CTD ’code’. Biosystems, 207, 104468.

30. Zaborowska, J., Egloff, S. and Murphy, S. (2016) The pol II CTD: new twists in the tail. Nature structural & molecular biology, 23, 771–777.

31. Jasnovidova, O. and Stefl, R. (2013) The CTD code of RNA polymerase II: a structural view. Wiley Interdiscip Rev RNA, 4, 1–16.

32. Whelan, M. and Pelchat, M. (2022) Role of RNA Polymerase II Promoter-Proximal Pausing in Viral Transcription. Viruses, 14, 2029.

33. Pflug, A., Guilligay, D., Reich, S. and Cusack, S. (2014) Structure of influenza A polymerase bound to the viral RNA promoter. Nature, 516, 355–360.

34. Wandzik, J.M., Kouba, T., Karuppasamy, M., Pflug, A., Drncova, P., Provaznik, J., Azevedo, N. and Cusack, S. (2020) A Structure-Based Model for the Complete Transcription Cycle of Influenza Polymerase. Cell, 181, 877–893 e821.

35. Fan, H., Walker, A.P., Carrique, L., Keown, J.R., Serna Martin, I., Karia, D., Sharps, J., Hengrung, N., Pardon, E., Steyaert, J. et al. (2019) Structures of influenza A virus RNA polymerase offer insight into viral genome replication. Nature, 573, 287–290.

36. Te Velthuis, A.J.W., Grimes, J.M. and Fodor, E. (2021) Structural insights into RNA polymerases of negative-sense RNA viruses. Nat Rev Microbiol, 19, 303–318.

37. Li, H., Wu, Y., Li, M., Guo, L., Gao, Y., Wang, Q., Zhang, J., Lai, Z., Zhang, X., Zhu, L. et al. (2023) An intermediate state allows influenza polymerase to switch smoothly between transcription and replication cycles. Nature structural & molecular biology, 30, 1183–1192.

38. Heidemann, M., Hintermair, C., Voss, K. and Eick, D. (2013) Dynamic phosphorylation patterns of RNA polymerase II CTD during transcription. Biochim Biophys Acta, 1829, 55–62.

39. Core, L. and Adelman, K. (2019) Promoter-proximal pausing of RNA polymerase II: a nexus of gene regulation. Genes Dev, 33, 960–982.

40. Fodor, E., Mingay, L.J., Crow, M., Deng, T. and Brownlee, G.G. (2003) A single amino acid mutation in the PA subunit of the influenza virus RNA polymerase promotes the generation of defective interfering RNAs. Journal of virology, 77, 5017–5020.

41. Carrique, L., Fan, H., Walker, A.P., Keown, J.R., Sharps, J., Staller, E., Barclay, W.S., Fodor, E. and Grimes, J.M. (2020) Host ANP32A mediates the assembly of the influenza virus replicase. Nature, 587, 638–643.

42. Staller, E., Sheppard, C.M., Neasham, P.J., Mistry, B., Peacock, T.P., Goldhill, D.H., Long, J.S. and Barclay, W.S. (2019) ANP32 Proteins Are Essential for Influenza Virus Replication in Human Cells. Journal of virology, 93.

43. Zhang, H., Zhang, Z., Wang, Y., Wang, M., Wang, X., Zhang, X., Ji, S., Du, C., Chen, H. and Wang, X. (2019) Fundamental Contribution and Host Range Determination of ANP32A and ANP32B in Influenza A Virus Polymerase Activity. Journal of virology, 93.

44. Boehning, M., Dugast-Darzacq, C., Rankovic, M., Hansen, A.S., Yu, T., Marie-Nelly, H., McSwiggen, D.T., Kokic, G., Dailey, G.M., Cramer, P. et al. (2018) RNA polymerase II clustering through carboxy-terminal domain phase separation. Nature structural & molecular biology, 25, 833–840.

45. Flores-Solis, D., Lushpinskaia, I.P., Polyansky, A.A., Changiarath, A., Boehning, M., Mirkovic, M., Walshe, J., Pietrek, L.M., Cramer, P., Stelzl, L.S. et al. (2023) Driving forces behind phase separation of the carboxy-terminal domain of RNA polymerase II. Nat Commun, 14, 5979.

46. Cho, W.K., Spille, J.H., Hecht, M., Lee, C., Li, C., Grube, V. and Cisse, II. (2018) Mediator and RNA polymerase II clusters associate in transcription-dependent condensates. Science, 361, 412–415.

47. Guo, Y.E., Manteiga, J.C., Henninger, J.E., Sabari, B.R., Dall’Agnese, A., Hannett, N.M., Spille, J.H., Afeyan, L.K., Zamudio, A.V., Shrinivas, K. et al. (2019) Pol II phosphorylation regulates a switch between transcriptional and splicing condensates. Nature, 572, 543–548.

48. Sabari, B.R., Dall’Agnese, A., Boija, A., Klein, I.A., Coffey, E.L., Shrinivas, K., Abraham, B.J., Hannett, N.M., Zamudio, A.V., Manteiga, J.C. et al. (2018) Coactivator condensation at super-enhancers links phase separation and gene control. Science, 361.

