## Supplementary Data for "Structural and functional characterisation of the interaction between the influenza A virus RNA polymerase and the CTD of host RNA Polymerase II"

| Peptide | Schematic and sequence |
|---|---|
| pS2 | <br>Y(pS)PTSPS Y(pS)PTSPS Y(pS)PTSPS Y(pS)PTSPS |
| pS5 | 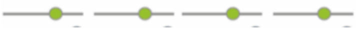<br>YSPT(pS)PS  YSPT(pS)PS  YSPT(pS)PS  YSPT(pS)PS |
| pS7 | <br>YSPTSP(pS) YSPTSP(pS) YSPTSP(pS) YSPTSP(pS) |
| pS(2/5).1* | 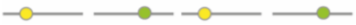<br>Y(pS)PTSPS  YSPT(pS)PS  Y(pS)PTSPS YSPT(pS)PS |
| pS(2/5).2* | <br>YSPT(pS)PS Y(pS)PTSPS YSPT(pS)PS Y(pS)PTSPS |
| pS(5/7).1* | 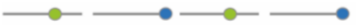<br>YSPT(pS)PS  YSPTSP(pS)  YSPT(pS)PS  YSPTSP(pS) |
| pS(5/7).2* | <br>YSPTSP(pS) YSPT(pS)PS YSPT(pS)PS YSPT(pS)PS |
| pS2 pS5** | <br>Y(pS)PT(pS)PS Y(pS)PT(pS)PS Y(pS)PT(pS)PS Y(pS)PT(pS)PS |
| pS5 pS7** | <br>YSPT(pS)P(pS) YSPT(pS)P(pS) YSPT(pS)P(pS) YSPT(pS)P(pS) |
| Scrambled<br>(control peptide) | PSSSTPSSYTPSPSSSPTSYSPYYTSPP |

**Supplementary Table 1.** Design of Pol II CTD mimic peptides with different phosphorylation states and simplified schematic to depict phosphorserines (pS) shown as coloured circles (pS2 (yellow), pS5 (green), pS7 (blue)). *Alternating phosphorylation of serine residues per heptad repeat. **Double phosphorylation of serine residues per heptad repeat.

| | 1918 vRNA 4rpt peptide<br>EMD-18945<br>PDB-8R60 | 1918 vRNA 4rpt peptide Ordered PB2<br>EMD-18947<br>PDB-8R65 |
|---|---|---|
| **Data collection** | | |
| Microscope | Titan Krios G3i (OPIC) | |
| Voltage (kV) | 300 | |
| Detector | Gatan K2 with EF | |
| Recording mode | Counting | |
| Magnification | 130,000 | |
| Movie/micrograph pixel size (Å) | 1.05 | |
| Dose rate (e-/px/sec) | 9.6 | |
| Number of frames per movie | 40 | |
| Movie exposure time (s) | 4.2 | |
| Total dose (e-/Å$^2$) | 40 | |
| Defocus range (um) | -1.2 to -2.5 | |
| **EM data processing** | | |
| Number of movies/micrographs | 7,921 | |
| Box size (px) | 256 | |
| Particle number (total) | 4,231,383 | |
| Particle number (used in final map) | 303,608 | 16,121 |
| Symmetry | C1 | |
| Map resolution (Å, FSC 0.143) | 3.23 | 4.23 |
| Map sharpening B-factor (Å$^2$) | 167.9 | 133.5 |
| **Model Building and Validation** | | **Rigid-body refinement only** |
| Initial model used | 7NHX | 7NHX/8R60 |
| Model composition | | |
| Non-hydrogen protein atoms | 27,951 | 35,542 |
| Protein residues | 1707 | 2196 |
| Ligands | 23 | 23 |
| B factors (Å$^2$) - mean | | |
| Protein | 77 | 317 |
| Nucleotide | 59 | 169 |
| RMSD from ideal | | |
| Bond length (Å) | 0.004 | 0.005 |
| Bond angles (°) | 0.501 | 0.621 |
| Validation | | |
| Molprobity score | 2.01 | 2.23 |
| Clashscore | 8.1 | 11.5 |
| FSC (0.5) model-vs-map | 3.5 | 7.6 |
| CC model-vs-map (masked) | 0.79 | 0.66 |
| Ramachandran plot | | |
| Favored (%) | 93.9 | 92.8 |
| Allowed (%) | 6 | 6.7 |
| Outliers (%) | 0.1 | 0.5 |

**Supplementary Table 2.** Cryo-EM data collection and model refinement statistics.

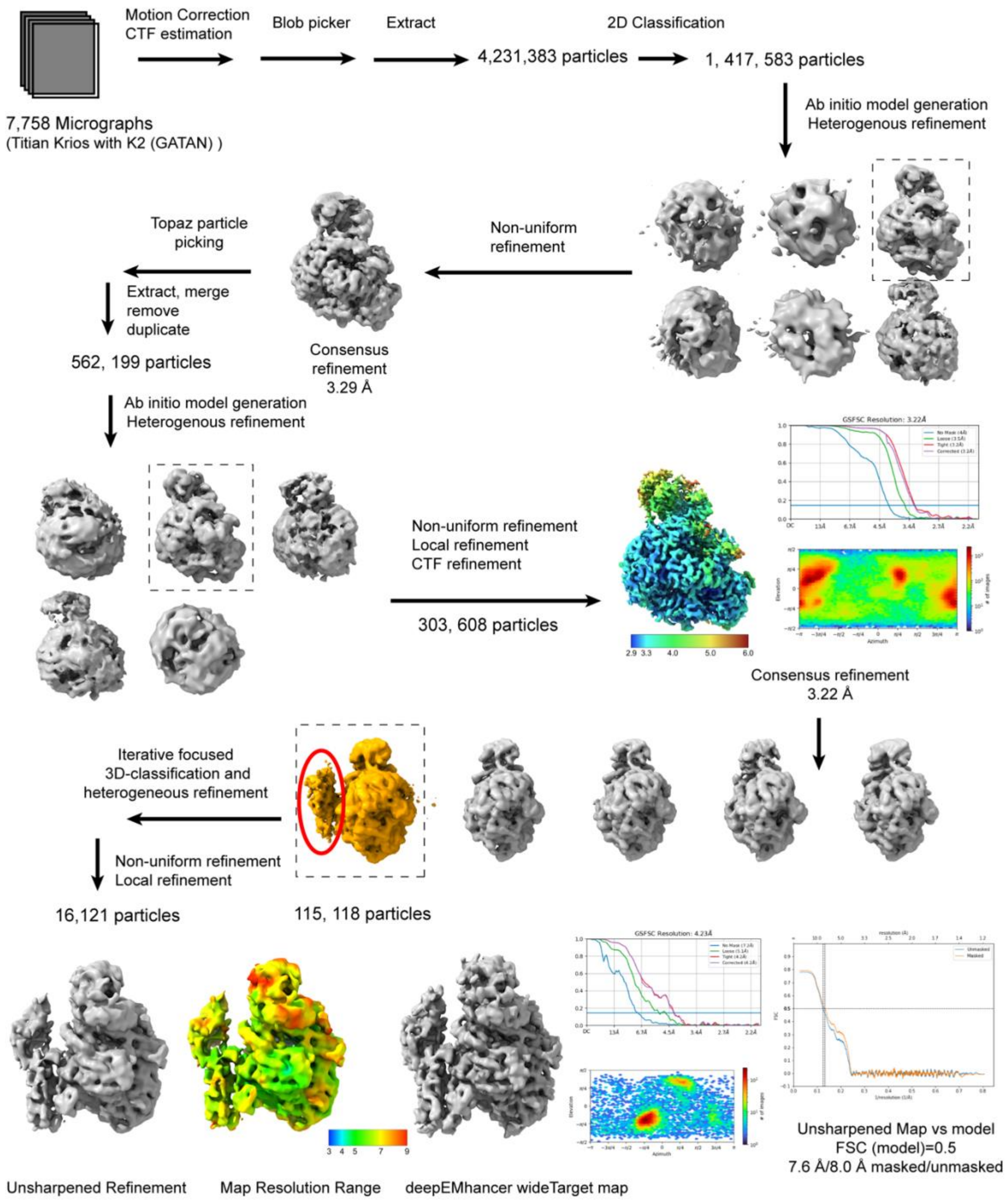

**Supplementary Figure 1.** Cryo-EM processing scheme.

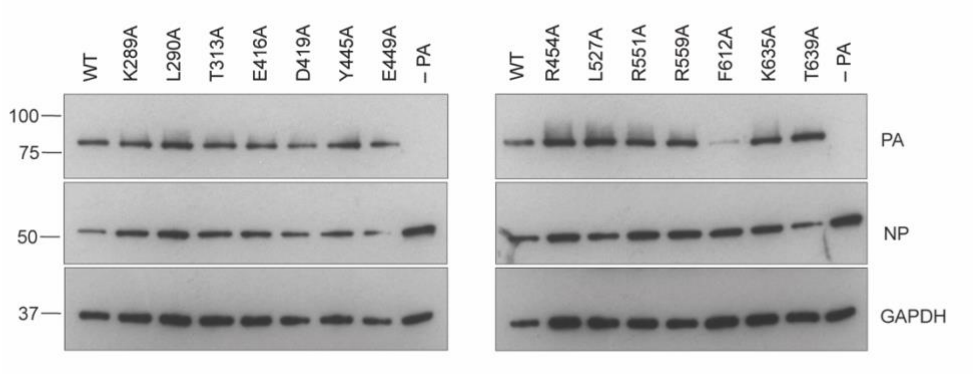

**Supplementary Figure 2.** Expression of the PA (WT and mutants) and NP of the 1918 pandemic H1N1 influenza A virus polymerase. HEK293T cells were transfected to express PB1, PB2 and PA (WT or mutant) polymerase subunits, NP, and segment 6 vRNA and cell lysates were analysed by SDS-PAGE and Western blotting with antibodies against PA, NP and GAPDH (loading control). A representative blot with size markers (kDa) is shown.

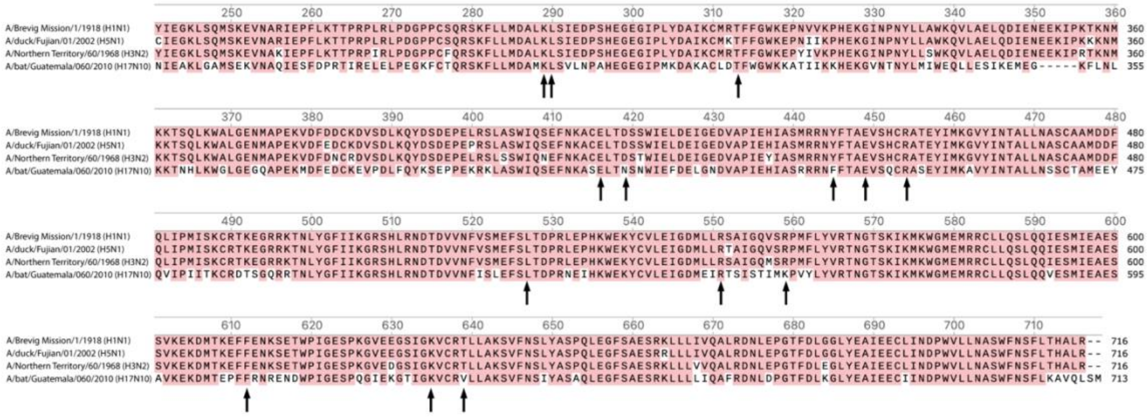

**Supplementary Figure 3.** Identification of conserved residues in the CTD-binding site of the influenza A virus PA subunit. Multiple sequence alignments of PA-C from A/Brevig Mission/1/1918 (H1N1), A/duck/Fujian/01/2002 (H5N1), A/Northern Territory/60/1968 (H3N2), and A/bat/Guatemala/060/2010 (H17N10) using Clustal-Omega in SnapGene. Identical residues are coloured in red. Black arrows depict conserved residues that were hypothesised to affect CTD binding based on our structural model of pS5 CTD-bound 1918 pandemic H1N1 influenza A virus polymerase.

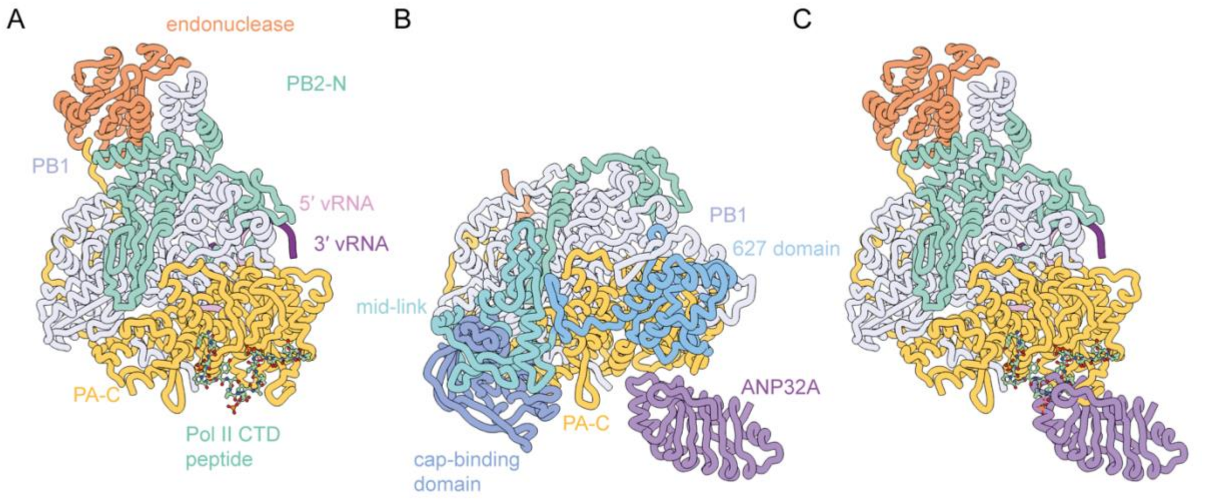

**Supplementary Figure 4.** Comparison of the Pol II CTD and ANP32 binding sites on the PA-C. (A) Influenza A virus polymerase bound to Pol II CTD peptide (PDB ID 8R60). (B) Influenza C virus polymerase in the encapsidating conformation bound to chicken ANP32A (PDB ID 6XZR). (C) Overlay of (A) and (B) indicating the overlap of the Pol II CTD and ANP32A binding sites on PA-C.
